# Functional and Structural Cerebellar-Behavior Relationships in Aging

**DOI:** 10.1101/2024.06.19.598916

**Authors:** Tracey H. Hicks, Thamires N. C. Magalhães, T. Bryan Jackson, Hannah K. Ballard, Ivan A. Herrejon, Jessica A. Bernard

## Abstract

Healthy aging is associated with deficits in cognitive performance and brain changes, including in the cerebellum. Yet, the precise link between cerebellar function/structure and cognition in aging remains poorly understood. We explored this relationship in 138 healthy adults (aged 35-86, 53% female) using resting-state functional connectivity MRI (fcMRI), cerebellar volume, and cognitive and motor assessments in an aging sample. We expected to find negative relationships between lobular volume for with age, and positive relationships between specific lobular volumes with motor and cognition respectively. We predicted lower cerebellar fcMRI to cortical networks and circuits with increased age. Behaviorally, we expected higher cerebello-frontal fcMRI cerebellar connectivity with association areas to correlate with better behavioral performance. Behavioral tasks broadly assessed attention, processing speed, working memory, episodic memory, and motor abilities. Correlations were conducted between cerebellar lobules I-IV, V, Crus I, Crus II, vermis VI and behavioral measures. We found lower volumes with increased age as well as bidirectional cerebellar connectivity relationships with increased age, consistent with literature on functional connectivity and network segregation in aging. Further, we revealed unique associations for both cerebellar structure and connectivity with comprehensive behavioral measures in a healthy aging population. Our findings underscore cerebellar involvement in behavior during aging.

## 1. Introduction

Healthy aging is associated with declines in cognitive and motor performance (1–4) as well as functional brain changes (5–9). To date, neuroimaging research has examined task-based functional activation and resting-state functional connectivity (FC) in aging adults as compared to young adults. Predominant theories of brain functional activation in the aging process include, 1) the hemispheric asymmetry reduction in older adults (HAROLD) characterized by less focal activation and increased bilateral hemispheric activation compared to younger adults during task performance (10), and 2) the compensation-related utilization of neural circuits hypothesis (CRUNCH) which suggests that the brain will recruit additional neural circuits to compensate for age-related declines in functioning (11). Both theories implicate more diffuse brain activation in aging as compared to young adults and align with FC findings of reduced network efficiency and increased integration of brain networks with greater age (5–7,9). Importantly, diffuse functional activation, lower network efficiency, and more integrated brain networks have been correlated with poorer cognitive performance (5,7,12,13). Older adults experiencing cognitive and motor deficits (e.g., memory problems, poor attention, reduced fine motor abilities, poor balance, etc.) often struggle with reduced independence, leading to a greater burden on the healthcare system and increased reliance on caregivers and family assistance (14,15). However, some older adults experience more deficits in healthy aging than others and the underlying causes of these differences remain poorly understood (16). A deeper understanding of cognitive and motor deficits in aging has the potential to inspire more precise interventions focused on alleviating the effects of behavioral and neurodegenerative-related deficits in later life.

In recent years, there has been an increased interest in understanding the cerebellum in aging. Anatomically, the whole cerebellum demonstrates decreased volume with increased age (17,18). Regional cerebellar volume also demonstrates significant age differences; however, results have varied with age in different lobular regions (19–24). For instance, Cui and colleagues (20) found lower volumes in lobules VI, X, Crus I, and Crus II (only 4 of the 12 regions examined) in older adults as compared to young adults; however, Han and colleagues (21) demonstrated significant negative relationships with age at baseline in 23 of the 28 lobules examined in older adults (ages ranged from 50-95 years old) with the exception of a positive relationship with right lobule IV and non-significant findings for left lobule IV, lobule VIIIA, and vermis VII. Further, a recent meta-analysis investigated cerebellar functional activation in young versus older adults (25). This study revealed that extant functional topography for the cerebellum was largely consistent between young and older adults (25). However, there was lower activation overlap for cognitive tasks and higher activation overlap in motor tasks for older as compared to young adults (25).

Finally, FC in the aging cerebellum has demonstrated mixed findings. He and colleagues (26) demonstrated intact cerebellar network segregation in aging adults (ages ranged from 19-80 years old) in a large sample, which contrasted with findings of lower network segregation with aging in several other networks such as the default mode (DMN) and fronto-parietal (FPN) networks. Dorsal and ventral cerebellar dentate to whole brain connectivity analyses in a large sample (*n* = 590, age range 18-88 years) has shown lower connectivity with increased age in adults (27), while lobular investigations have also shown broad patterns of lower connectivity in older relative to young adults (28). Alternatively, Seidler and colleagues (29) found that in the context of motor function, older individuals exhibited increased connectivity between cerebellar lobule VIII and the putamen, and decreased connectivity between lobule V and cortical regions. These studies suggest that patterns of FC vary within regions of the cerebellum, with age, or depending on the task. Notably, clear patterns of cerebellar connectivity in aging have yet to be established and more work is needed in this area.

Across a growing literature, alterations in cerebellar structure and functional networks have been implicated in the cognitive and motor deficits observed in older adults (19,20,30–34). Evaluating cerebellar impacts on behavior across adulthood and aging is less explored, although it is clear that the cerebellum contributes to both motor and cognitive behaviors (25,35–38). Specifically, functional topography of the cerebellum derived from task-based activation has been useful in mapping cognitive, motor, and emotional contributions (25,37–40). While cerebellar functional topography is not confined by anatomical boundaries, the differences seen in functional activation across regions are important for increasing our understanding of behavioral deficits in aging, regional insults, or pathological neurodegenerative processes. The work to date investigating cerebellar metrics in advanced age have linked both structure and resting state networks to motor and cognitive performance in older adults (28,30,31), though the samples here were relatively small and it was a limited behavioral battery. While Koppelmans et al. (32,41) looked at a much larger sample and also demonstrated associations with behavior, the age range was limited to those over age 65 and only considered cerebellar structure. Thus, while this work has provided a strong foundation of understanding with respect to age differences in cerebellar metrics and behavior, this work has focused almost exclusively on comparisons between young and older adults. Understanding these relationships across adulthood, with the inclusion of middle age where subtle changes in brain and behavior are likely to begin, is critical. Further, resting state lobular networks have only been explored in a small sample and have been given limited consideration with respect to behavior (28). The cerebellum has been conceptualized as key scaffolding that is critical for behavioral performance and further influences cortical function (42), and as such, understanding this scaffolding across the adult lifespan stands to provide needed insights into age-related behavioral decline.

Here, we investigated the cerebello-cortical FC and structural volume of cerebellar lobules in association with an extensive battery of cognitive and motor behaviors. It is well accepted that the cerebellum communicates with the cortex via closed-loop circuits through the thalamus (43–45), that in turn relate to the cerebellar functional topography. That is, work using task-based neuroimaging has shown that distinct regions of the cerebellum are associated with different behavioral domains (37,38,46). This organization is, not surprisingly, consistent with known structural connections to the motor and prefrontal cortices, and also follows patterns of cortical functional network presence in the cerebellum (37). This highlights the necessity of taking a more nuanced and focused approach to investigating cerebellar-behavior relationships. While extant functional topography and network parcellations do not exclusively follow lobular anatomical boundaries (28,37,47), we have narrowed our investigation here to a subset of cerebellar lobules in a hypothesis driven way. This approach also allows us to limit (at least to a degree) concerns about multiple comparisons. Here we are focused on lobules I-IV, V, Crus I, Crus II, and vermis VI. Functionally, lobules I-IV and V have largely been associated with motor function (37,39,46), whereas Crus I and II are implicated in cognitive behaviors (37,38,46) in functional activation studies, and share networks with frontal and association areas of the cortex (28,47,48) in FC studies. Structurally, greater Crus I volume has been associated with higher-level cognitive processing (30), greater left posterior cerebellar volume (including lobules VIIb, VIIIa, VIIIb, IX, X, and Crus I and II) was linked to working memory accuracy in older adults, (30), and greater volumes in bilateral Crus I, lobule VI, and right lobule IV were associated with better memory performance in aging adults (20). Vermis VI allows for broader exploration of midline cerebellar contributions to behavior. This study examined lobular regions via volumetric structure and lobular seed-based connectivity to examine specific cerebellar regions that have been linked to changes with aging and behavioral performance.

Few studies have examined functional networks between cerebellar regions of interest and the cortex alongside behavioral performance data. Namely, one study revealed age-related differences in cerebellar to basal ganglia connectivity were associated with behavioral performance (i.e., global cognition screener and balance confidence) in older adults (31). Another study by Bernard and colleagues (28) showed FC associations between Crus II and the posterior cingulate cortex in relation to working memory in older adults. Here, we intend to increase the depth of literature for cerebello-cortical connectivity in the context of more diverse cognitive assessments and more expansive evaluation of cortical regions. Cortically, there have been consistent findings showing decreases in FC within the DMN in healthy aging (6,26,49,50) which has been linked to poorer behavioral performance (see review 51). It has been theorized that Crus II may play a role in the DMN (28). Thus, cortical FC relationships between cerebellar regions and aspects of the DMN are of special interest in our study. Additionally, there have been consistent findings of Crus I and Crus II connectivity to the prefrontal cortex (47,48,52), implicating connectivity between these regions as a possibly important connection for cerebello-cortical networks in aging and behavior. We hypothesized that age would negatively correlate with volume across ROIs. Further, we predicted that cerebello-cortical FC would be lower across lobules and particularly with regions of the DMN, with increased age. We further predicted that stronger Crus I and Crus II FC to prefrontal and association regions would be associated with better cognitive performance. Similarly, we expected positive correlations between Crus I and Crus II volume and cognitive performance. Lobules I-IV and V volumes were expected to display positive relationships with motor measures and an integrated cognitive-motor task. Further, we expected higher FC between motor-related lobules (lobules I-IV and V) and motor cortical regions to correlate with better motor performances. Our lobular approach stands to bolster extant functional and structural findings in the cerebellum. By exploring these associations, we seek to enhance our understanding of age-related cognitive and motor decline.

## Methods

### 1.1. Study sample

Participants (total *n*=175) were enrolled as part of a larger study on aging. All participants underwent a battery of cognitive and motor tasks during this assessment described in detail below. After the behavioral visit the participants returned for a magnetic resonance imaging (MRI) session approximately two weeks later. However, due to unexpected delays related to the Covid-19 pandemic, the time between the two sessions (39.0 days ± 21.4 days) varied between participants. For the context of this study, we focused on specific aspects of cognitive and motor performance relationships with brain imaging data.

Exclusion criteria were history of neurological disease, stroke, or formal diagnosis of psychiatric illness (e.g., depression or anxiety), contraindications for the brain imaging environment, diagnosis of mild cognitive impairment or dementia, and use of hormone therapy (HTh) or hormonal contraceptives (intrauterine device (IUD), possible use of continuous birth control (oral), and no history of hysterectomy in the past 10 years. For our analyses here we focused only on those with available neuroimaging and cognitive performance data. Thus, our final sample included 138 participants, 54% female (74 females (*M* = 56.73, SD = 12.53) and 64 males (*M* = 56.58, SD = 12.53)). Ages ranged from 35 to 86 years. Regarding ethnicity, our sample was 88.4% Caucasian and 11.6% Hispanic/Latino. The distribution of race is listed below (Table 1). Demographic details are presented for generalizability purposes and we recognize that these are social and political categories given meaning by social, historical, and political forces (53). Race and ethnicity will not be evaluated separately in this study, as information about socio-economic status is better indicated for causal inferences in neuroimaging (53) and that information was not collected in this study.

**Table 1.** Demographic means for the full sample are listed below with standard deviations in parentheses.

| Demographics |  |
| --- | --- |
| Age (years) | 57(13.31) |
| Sex F(M) | 74(64) |
| African American (%) | 2% |
| Asian (%) | 7% |
| Caucasian (%) | 86% |
| Multiracial (%) | 4% |
| Native American (%) | 1% |

**Table 2.** Means for the behavioral tasks are listed below with standard deviations in parentheses.

| Behavioral Task Performance |  |  |
| --- | --- | --- |
| Variable | Value | Mean (SD) |
| Comp1 | Sum of z-scored measures | -0.040<br>(3.066) |
| Comp2 | Sum of percent correct across measures | 1.641<br>(0.349) |
| Stroop Effect Reaction Time | Difference between average incongruent and congruent item responses in milliseconds | 101.659<br>(77.488) |
| Sequence Learning | Learning slope coefficient across consecutive blocks | 0.011<br>(0.026) |
| Purdue Pegboard Assembly | Z-scored, a higher score indicates better performance | -0.144<br>(0.962) |
| Motor Composite | Sum of z-scored measures | 0.819<br>(4.496) |

All study procedures were approved by the Institutional Review Board at Texas A&M University, and written informed consent was obtained from each participant prior to initiating any data collection.

### 1.2. Behavioral Testing

Participants completed a battery of cognitive and motor tasks to quantify attention, processing speed, working memory, episodic memory, executive functioning, grip strength, fine motor abilities, and cognitive-motor integration. A commonly used screening tool that assesses global cognitive functioning, the Montreal Cognitive Assessment (MoCA) was also included (54). The Wechsler Adult Intelligence Scale, 4th Edition (WAIS-IV) Digit Span and Letter-Number Sequencing subtests (55) broadly assessed simple attention and working memory. The WAIS-IV Coding subtest was used to gauge attention and processing speed. The Wechsler Memory Scale, 4^th^ Edition (WMS-IV) Symbol Span subtest was used to assess visual-spatial working memory (56).

Episodic memory was assessed via the Shopping List Memory Task (57) which was administered on a computer. Words of 30 items resembling those commonly seen on a shopping list were presented to the participant one at a time for 14 seconds each with a 0.5 second fixation cross in between words. Memory for the items was assessed approximately 20 minutes later via recognition discrimination for the original list via percentage of items answered correctly.

#### 1.2.1. The Stroop Task

The Stroop Task (58) was adapted to computer format and was used to gauge executive function. Both congruent and incongruent items were displayed for 2.6 seconds each. There were 10 blocks with untimed (self-paced) breaks in between each block. This study specifically examined the Stroop effect as an outcome. Mathematically, the score was the difference in mean accuracy between congruent (e.g., the word “blue” in blue font) and incongruent (e.g., the word “blue” in red font) items (incongruent – congruent). We also examined the Stroop effect in the context of reaction time differences between congruent and incongruent items in which a higher score indicated a slower reaction time on incongruent trials, score of 0 indicated performing the same (on average) for both congruent and incongruent trials, and a negative score indicated a faster reaction time on incongruent trials. Those with incomplete data were also excluded from calculations for this task (n = 1; 9/10 blocks incomplete).

#### 1.2.2. Sequence Learning Paradigm

An explicit sequence learning paradigm was based on a task created by Bo and colleagues (59). This task assessed learning of motor behaviors and working memory. The task consisted of 6 random blocks with 18 items each interspersed with 9 sequence blocks with 36 items each. In sequence blocks, participants learned a 12-element sequence shown 1 second per element and repeated this sequence 4 times in each block.

In the random blocks, participants learned a different sequence each time. For both blocks, they were shown a sequence of filled squares and asked to repeat it in its entirety. We chose to quantify learning by calculating the slope of task accuracy (labeled as sequence learning accuracy in the results section) from Block 1 to Block 9. Conceptually, we would expect a positive slope for accuracy if learning had occurred.

Cognitive composite variables were created based on the format of task administration (i.e., paper and pencil type, computer-based). Raw scores from each cognitive task were z-scored across the sample. These z-scores were then added together across Cognitive composite 1 (Comp1) to create composite cognitive scores for each participant. Comp1 included Coding, Digit Span, Symbol Span, and Letter-Number Sequencing, and broadly represented attention, processing speed, and working memory. Cognitive composite 2 (Comp2) included accuracy-based measures (sum of the percent correct) on each of the following computer-based tasks: shopping list memory, Stroop effect (accuracy incongruent trials – accuracy congruent trials), and overall performance accuracy on a sequence learning task. Comp2 broadly represents episodic memory, executive function, and motor learning across these computer-based tasks.

Motor function was assessed via a motor composite variable using the Purdue Pegboard Task (60) and a bilateral assessment of grip strength which quantifies fine and gross motor upper limb function, respectively. The Purdue Pegboard is comprised of four subtests: dominant hand, non-dominant hand, both hands simultaneously, and assembly task. Three of the subtests require simple fine motor skills of placing a peg into a groove down one or two lines of grooves depending on the task. Lastly, there is the assembly subtest where participants “assemble multiple components into a unit, which is made by placing a peg in a hole (dominant hand), placing a washer over the peg (nondominant hand), then a small cylindrical collar (dominant hand), followed by a second washer on top (non-dominant hand)” (Wilkes et al., 2023). This requires fine motor function as well as sequencing skills and bimanual coordination. Grip strength was measured using a handheld dynamometer. Participants were asked to grip the handle as hard as they could with their dominant hand for 5 seconds. Like with the cognitive tasks, our motor composite was comprised of the sum z-scored values from the following subtests: Purdue Pegboard dominant, non-dominant, both hands, and assembly tasks, average right-handed grip strength, and average left-handed grip strength. The Purdue Pegboard Assembly was also assessed separately due to the integration of cognitive and motor demands of the task which is particularly relevant to our investigation of the cerebellum and cerebello-cortical networks.

### 1.3. Imaging acquisition

The imaging sections (section 1.3 through 2.3.1.2.) utilize standardized text to ensure consistency and reproducibility in the research protocol. These standardized descriptions can also be found in other publications from our laboratory (62,63) and align with current best practices in the field, providing a clear and detailed framework for the procedures undertaken.

Participants underwent structural and resting-state MRI using a Siemens Magnetom Verio 3.0 Tesla scanner and a 32-channel head coil. For structural MRI, we collected a high-resolution T1-weighted 3D magnetization prepared rapid gradient multi-echo (MPRAGE) scan (repetition time (TR) = 2400 ms; acquisition time = 7 minutes; voxel size = 0.8 mm^3^) and a high-resolution T2-weighted scan (TR = 3200 ms; acquisition time = 5.5 minutes; voxel size = 0.8 mm^3^), each with a multiband acceleration factor of 2. For resting-state imaging, we administered four blood-oxygen level dependent (BOLD) functional connectivity (fcMRI) scans with the following parameters: multiband factor of 8, 488 volumes, TR of 720 ms, and 2.5 mm^3^ voxels. Each fcMRI scan was 6 minutes in length for a total of 24 minutes of resting-state imaging, and scans were acquired with alternating phase encoding directions (i.e., two anterior to posterior scans and two posteriors to anterior scans). During the fcMRI scans, participants were asked to lie still with their eyes open while fixating on a central cross. In total, the acquisition of images took about 45 minutes, including a 1.5-minute localizer.

Scanning protocols were adapted from the multiband sequences developed by the Human Connectome Project (HCP) (64) and the Center for Magnetic Resonance Research at the University of Minnesota to facilitate future data sharing and reproducibility.

#### 1.3.1. Imaging processing

##### 1.3.1.1. Pre-processing

The images underwent several preprocessing steps to prepare them for further analysis. Initially, they were converted from DICOM to NIFTI format and organized following the Brain Imaging Data Structure (BIDS, version 1.6.0) using the bidskit docker container (version 2021.6.14, https://github.com/jmtyszka/bidskit). Afterward, a single volume was extracted from two oppositely coded BOLD images to estimate B0 field maps using the split tool from the FMRIB Software Library (FSL) package (Jenkinson et al., 2012). Subsequently, the anatomical and functional images were preprocessed using fMRIPrep (version 20.2.3; for detailed methods, see https://fmriprep.org/), which includes automated procedures to align the functional volume with the anatomical image, correct for motion, correct field map distortions, segment the anatomical image into distinct tissues (e.g., gray matter, white matter, cerebrospinal fluid), remove the skull from the anatomical image, normalize the data to a common space, align motion-corrected functional volumes with the normalized anatomical image, and apply spatial smoothing.

##### 1.3.1.2. Functional analysis

Following preprocessing with fMRIPrep, the subsequent analyses were conducted using the CONN toolbox (version 21a) (65). This involved additional processing to eliminate noise and artifacts and enhance data quality. Denoising in CONN typically comprises several stages, such as removing motion signals and regressing out confounding signals (e.g., signals from ventricles, white matter, and global signals). Motion information from fMRIPrep was utilized in CONN for this purpose. A 0.008-0.099 Hz bandpass filter was applied to eliminate high-frequency noise. The denoising step is crucial for enhancing the quality of FC data by minimizing artifacts and enhancing the ability to detect genuine FC patterns in the brain.

In the first level analyses, we included the following cerebellar regions of interest (ROIs): lobules I-IV, V, Crus I, Crus II, and vermis VI. Lobular seeds for the ROIs were originally created based on the regional specifications from SUIT atlas (66) by Bernard and colleagues and later applied to our current sample (28). The SUIT atlas was created for anatomically-specific cerebellar investigations (66,67). Lobules I, II, III, and IV are combined in the SUIT atlas so they were investigated together. The investigation was limited to the right hemisphere for Crus I, Crus II, and lobule V when looking at hemispheric lobules, to mitigate multiple comparisons and lobules were selected based on our a priori hypotheses with respect to cerebellar-behavior associations, as described in the Introduction. Whole brain seed-to-voxel analyses were conducted at the group level, looking at both connectivity patterns with respect to age, but also as they predict behavior, using correlation approaches. We employed standard settings for cluster-based inferences using parametric statistics based on random field theory. We used an initial voxel threshold at p<.001 along with a cluster threshold set at p < .05, with a false discovery rate (FDR) correction.

##### 1.3.1.3. Structural processing

As described earlier, we began by converting high-resolution T1-weighted 3D images from DICOM to NIFTI format. Subsequently, we utilized FastSurfer software (https://github.com/Deep-MI/FastSurfer) in conjunction with the latest version of FreeSurfer (version 7.4.1) (https://github.com/freesurfer/freesurfer) (68,69), and CerebNet (70) into our structural analysis. FastSurfer, is specifically designed to automatically segment cortical gray matter and calculate key morphometric measurements including cortical thickness, surface area, and volume. The image analysis workflow involves pre-processing, corrects signal intensity inconsistencies caused by polarization fields, and involves removing the skull and non-brain tissues from MRI images. CerebNet is specialized in cerebellar analysis and its subregions, offering tools for cerebellar segmentation, surface reconstruction, and morphometric measurements. It is a fully automated and extensively validated deep-learning method for cerebellar lobular segmentation (70). CerebNet combines FastSurferCNN, a UNet-based 2.5D segmentation network, with advanced data augmentation techniques including realistic non-linear deformations. This approach enhances anatomical variability and eliminates the need for additional preprocessing steps like spatial normalization or bias field correction. CerebNet has demonstrated exceptional accuracy, surpassing current state-of-the-art approaches in cerebellar segmentation (70). Regions of interest (ROIs) included lobules I-IV, right Crus I, right Crus II, right lobule V, and vermis VI. Total intracranial volume (TIV) was used to correct for overall head size for standardization of ROI volume in later analyses and TIV values were extracted from Freesurfer output.

### 1.4. Statistical analyses

For statistical analyses of the structural results, we used IBM SPSS software (version 29, SPSS Inc., Chicago, IL, USA). Structural analyses were conducted concurrently with connectivity analyses to explore potential parallel relationships between volume in the cerebellar ROIs and the selected variables (age, Comp1, Comp2, Stroop effect for reaction time, Sequence learning, Purdue Pegboard Assembly, and motor composite).

We first sought to ascertain relationships via correlations between age and cortical FC, as well as partial correlations for age and TIV-corrected volume for each ROI (i.e., lobules I-IV, V, Crus I, Crus II, and vermis VI) within our sample. To determine brain-behavior performance relationships, the variables Comp1, Comp2, Stroop effect for reaction time, Sequence learning, Motor Composite, and Purdue Pegboard Assembly were each correlated with cortical FC for each ROI (respectively). We used the same statistical cut-offs and multiple comparisons corrections outlined above. Separate partial correlations between the aforementioned variables and TIV-corrected ROI volumes were conducted. All analyses were corrected for multiple comparisons using an FDR correction.

## 1. Results

### Age is Associated with Lobular Connectivity and Volume

Age was significantly correlated with connectivity in lobules I-IV, V, Crus I, and Crus II (Figure 1) but was not significant in vermis VI (Table 3). Increased age correlated significantly with higher connectivity between lobules I-IV and the caudate (Figure 1B) and higher connectivity between Crus II and the anterior cingulum and lingual gyrus (Figure1A). Higher connectivity between Crus I and the lingual gyrus also significantly correlated with increased age (Figure 1D). Greater connectivity was significantly correlated with increased age between lobule V and the caudate, superior frontal gyrus, thalamus, and olfactory cortex (not visualized). However, there were also findings in the opposite direction. Increased age was significantly correlated with lower connectivity between Crus II and the middle temporal gyrus, lobule VI, and superior and inferior parietal gyrus (Figure 1A), and with lower connectivity between lobule V and the precuneus and calcarine cortex (Figure 1C). This is consistent with mixed findings seen in prior work in this area (26,27,29).

**Figure 1.**
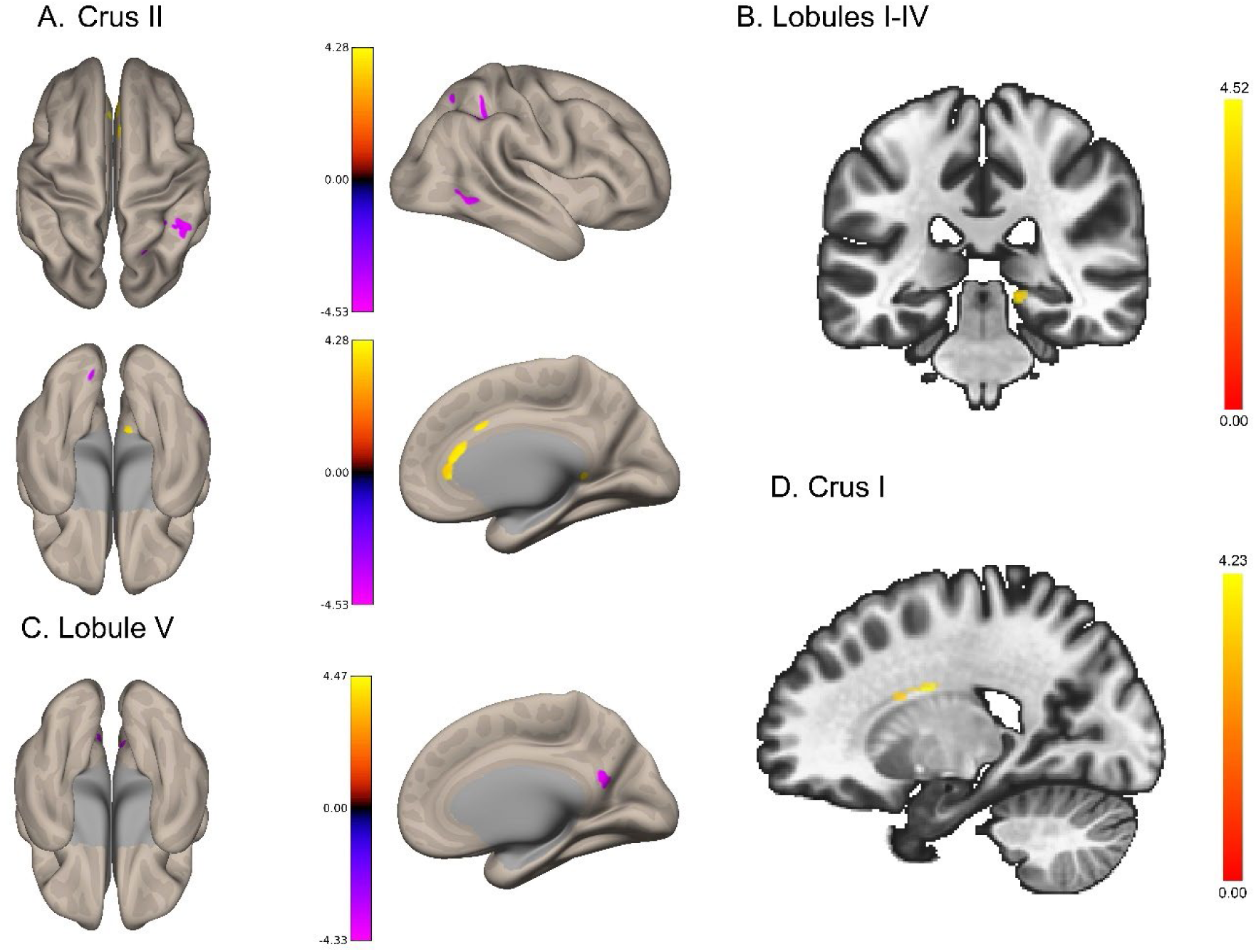
Patterns of cortical functional connectivity (FC) with ROIs. **A**. purple represents lower FC between Crus II and the inferior & superior parietal gyrus, and middle temporal gyrus (lobule VI not visualized) with increased age; while yellow displays greater FC between Crus II and both the anterior cingulum and lingual gyrus with increased age. **B**. yellow represents greater FC between the caudate and lobules I-IV with age. **C**. purple displays lower FC between lobule V and both the calcarine cortex and precuneus with greater age (ROIs with greater FC are listed in Table 1 and not visualized). **D**. yellow represents greater FC between Crus I and the lingual gyrus with increased age.

**Table 3.** Coordinates of cerebellar regions showing significant correlations in cortical FC with age.

| Functional Connectivity Correlations with Age |  |  |  |  |  |  |
| --- | --- | --- | --- | --- | --- | --- |
| Region | X | Y | Z | Cluster Size | T(136) | pFDR |
| **Lobules I-IV** |  |  |  |  |  |  |
| Caudate | 20 | -4 | 28 | 25 | 5.18 | 0.0059 |
| **Crus I** |  |  |  |  |  |  |
| Lingual gyrus | 16 | -32 | -10 | 29 | 5.27 | 0.0029 |
| **Crus II** |  |  |  |  |  |  |
| Inferior Parietal gyrus | 46 | -44 | 50 | 42 | -5.69 | 0.0002 |
| Middle Temporal gyrus | 60 | -58 | 0 | 33 | -6.22 | 0.0007 |
| Anterior Cingulum | 0 | 24 | 20 | 30 | 6.38 | 0.0009 |
| Anterior Cingulum | 2 | 36 | 6 | 23 | 6.34 | 0.0043 |
| Lingual gyrus | 8 | -38 | 2 | 22 | 4.66 | 0.0046 |
| Anterior Cingulum | 0 | 22 | 30 | 18 | 4.68 | 0.0119 |
| Superior Parietal gyrus | 28 | -74 | 56 | 14 | -5.36 | 0.0307 |
| Lobule VI | -8 | -78 | -14 | 14 | -4.88 | 0.0307 |
| **Lobule V** |  |  |  |  |  |  |
| Caudate | -20 | -2 | 28 | 42 | 5.94 | 0.0002 |
| Superior Frontal gyrus | 26 | 38 | 8 | 29 | 6.06 | 0.0019 |
| Thalamus | 0 | -26 | 10 | 28 | 5.40 | 0.0019 |
| Calcarine cortex | -4 | -68 | 22 | 26 | -5.52 | 0.0024 |
| Caudate | 20 | 4 | 26 | 24 | 5.15 | 0.0033 |
| Precuneus | 6 | -56 | 20 | 22 | -5.84 | 0.0048 |
| Olfactory cortex<br>(Caudate) | 2 | 16 | 2 | 17 | 5.76 | 0.0178 |
| Caudate | -22 | -18 | 30 | 15 | 4.88 | 0.0291 |

With respect to structure, partial correlations between ROI volume, correcting for TIV, and age revealed significant negative relationships across all ROIs (see Table 4 for statistics, and Figure 2 for a visualization). With increasing age, volume was significantly smaller across all ROIs that were investigated.

**Figure 2.**
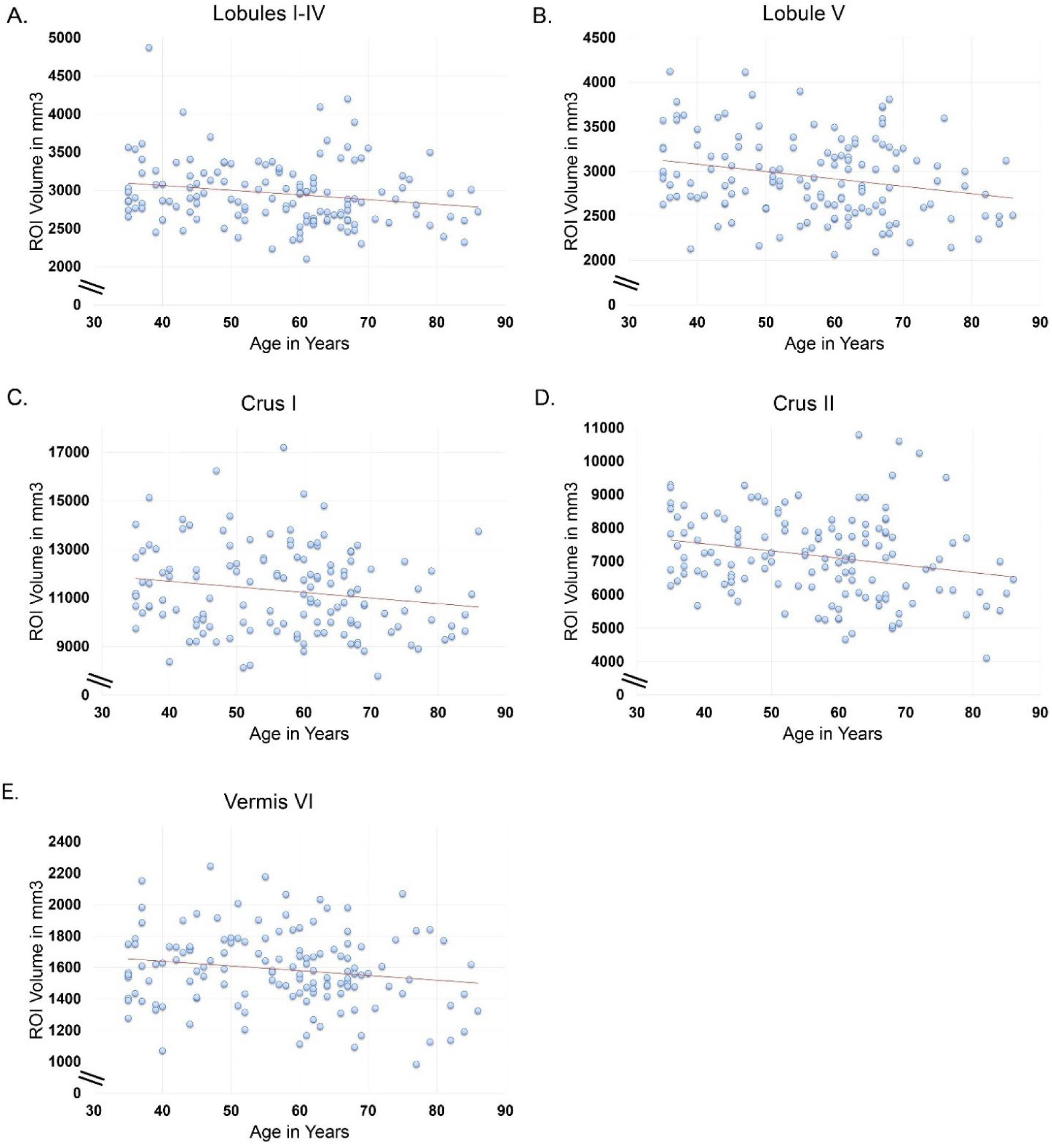
Negative correlations between age and structural volume across ROIs. See Table 3 for statistical values. These figures display rote correlations that do not account for TIV and are solely presented for visualization purposes.

**Figure 3.**
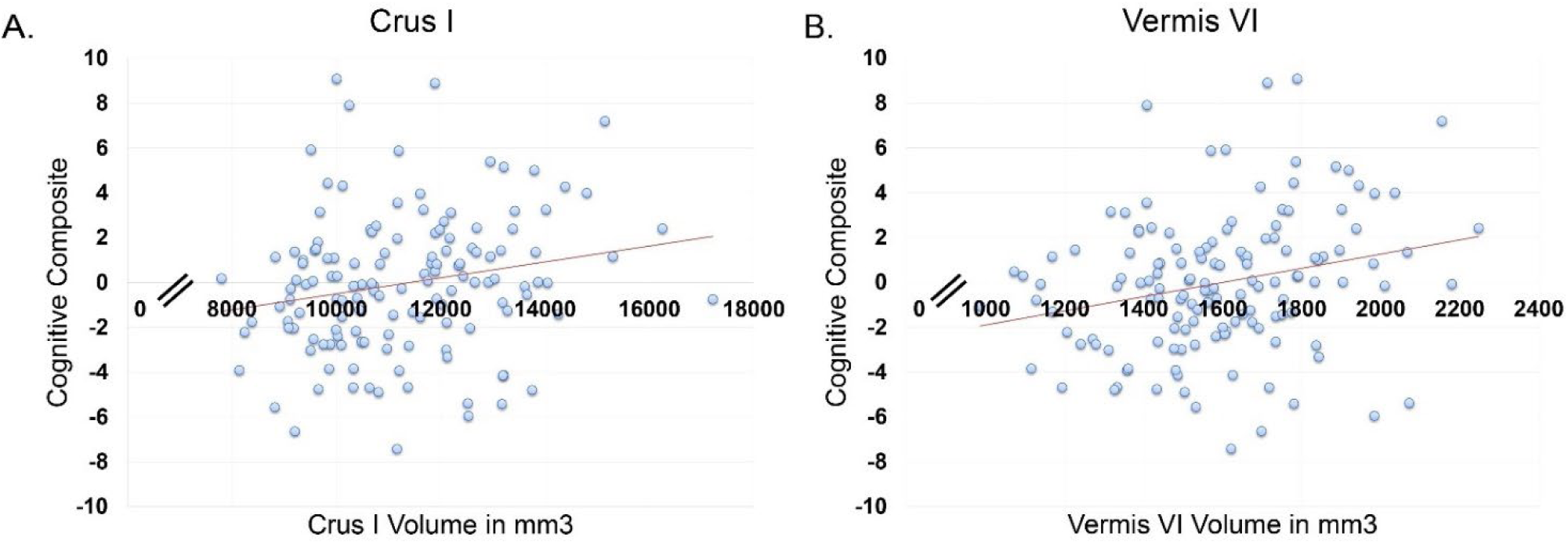
**A**. Crus I volume shows a positive correlation with the cognitive composite (Comp1). **B**. Vermis VI also displays a positive correlation with Comp1. These figures display rote correlations that do not account for TIV and are solely presented for visualization purposes.

**Table 4.** Cerebellar regional volumes show significant negative correlations with age.

| **ROI** | ***r*** | **df** | **pFDR** |
| --- | --- | --- | --- |
| Lobules I-IV | -0.267 | 135 | 0.005 |
| Crus I | -0.231 | 135 | 0.009 |
| Crus II | -0.253 | 135 | 0.005 |
| Lobule V | -0.323 | 135 | <0.001 |
| Vermis VI | -0.201 | 135 | 0.019 |

### Behavior is Associated with Lobular Connectivity and Volume

There were no significant associations between Comp 1 (Digit Span, Symbol Span, Letter-Number Sequencing, and Digit-Symbol Coding) and lobular connectivity (all p_FDR_>0.05). Partial correlations between ROI volume, controlling for TIV, and Comp1 revealed significant positive relationships with Crus I (*r* (135) = 0.220, p_FDR_ =0.010; Figure 2A) and vermis VI (*r* (135) = 0.260, p_FDR_ =0.01; Figure 2B). Larger volume in these regions predicted better performance. Partial correlations were not significant between Comp1 and lobules I-IV, lobule V, and Crus II (p_FDR_ > 0.05).

With respect to Comp2 (Shopping List, Sequencing, and Stroop accuracy) and lobular connectivity analyses, better performance on Comp2 was significantly correlated to stronger connectivity between Crus I and the superior parietal gyrus (T(102) = 5.34, p_FDR_ = 0.0772; X,Y,Z: -28, -56, 46; cluster size 17). We did not find significant associations between Comp2 and connectivity of the remaining ROIs (p_FDR_ > 0.05). Partial correlations between TIV, ROI volume, and Comp2 did not reveal significant relationships across ROIs (p_FDR_ > 0.05).

The Stroop effect for reaction time (average reaction time for incongruent – congruent trials) was also examined in the context of ROI to whole brain connectivity and lobular volume. A higher value for the Stroop effect reaction time variable indicated a slower response on incongruent trials as compared to congruent trials performance.

Thus, a higher value for this variable indicates a larger Stroop effect. A larger Stroop effect was significantly correlated with lower connectivity between Crus II and cerebellum VI (Figure 4B, Table 5). Lobule V demonstrated both positive and negative correlations with the Stroop effect. Specifically, a stronger Stroop effect was correlated with lower connectivity between lobule V and both the middle frontal gyrus and inferior parietal gyrus. Higher connectivity between lobule V and the postcentral gyrus (right sensory association) was positively correlated with a stronger Stroop effect (Figure 4C). Lastly, lower connectivity between vermis VI and regions of the middle frontal gyrus were significantly correlated with a stronger Stroop effect (Figure 4A). Partial correlations were not significant between TIV, ROI volume, and the Stroop effect for reaction time across regions (p_FDR_ > 0.05).

**Figure 4.**
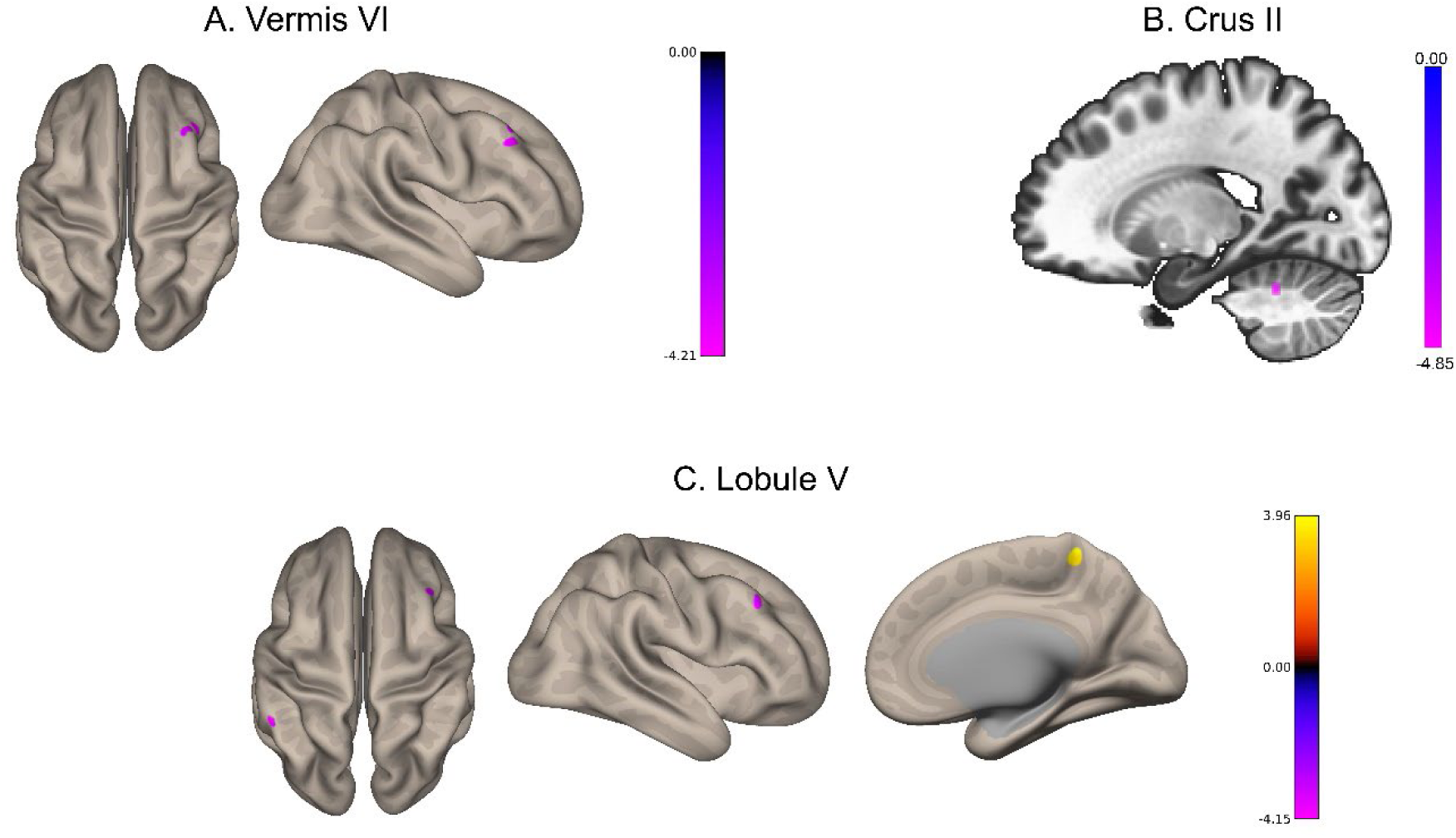
Bidirectional cortical functional connectivity (FC) is shown in ROIs with a stronger Stroop effect reaction time. **A**. purple represents lower FC between vermis VI and the middle frontal gyrus with a stronger Stroop effect. B. purple represents lower FC between Crus II and cerebellum VI with a stronger Stroop effect. C. purple represents lower FC between both the middle frontal gyrus and inferior parietal gyrus and lobule V with a stronger Stroop effect; yellow represents greater FC between lobule V and and the postcentral gyrus with a stronger Stroop effect.

**Table 5.** Coordinates of regions showing significant correlations between cerebellar connectivity and the Stroop Effect for reaction time.

| Functional Connectivity Correlations<br>with Stroop Effect for Reaction Time |  |  |  |  |  |  |
| --- | --- | --- | --- | --- | --- | --- |
| Region | X | Y | Z | Cluster<br>Size | T(127) | pFDR |
| **Crus II** |  |  |  |  |  |  |
| Cerebellum VI | -18 | -52 | -30 | 18 | -5.98 | 0.0265 |
| **Lobule V** |  |  |  |  |  |  |
| Middle frontal gyrus | 46 | 28 | 40 | 26 | -4.55 | 0.0016 |
| Postcentral gyrus | 16 | -42 | 66 | 20 | 4.38 | 0.0046 |
| Inferior parietal gyrus | -52 | -42 | 48 | 13 | -5.64 | 0.0316 |
| **Vermis VI** |  |  |  |  |  |  |
| Middle frontal gyrus | 46 | 32 | 36 | 21 | -5.66 | 0.0130 |
| Middle frontal gyrus | 38 | 32 | 48 | 15 | -5.55 | 0.0435 |

To evaluate associations with learning and motor abilities, accuracy and reaction time across sequencing trials were correlated with ROI connectivity. Learning was calculated via the percent accuracy slope across 9 consecutive trials. Here a higher score indicates improvement in integrated cognitive-motor learning across blocks. Learning performance correlations with ROI connectivity were significant for Crus II and lobule V (Figure 5, Table 6). Specifically, greater learning was correlated with lower connectivity between Crus II and the precentral gyrus (Figure 5B; Table 6). Further, stronger connectivity between lobule V and regions of the inferior parietal gyrus were significantly correlated with a better learning performance on the Sequencing task (Figure 5A). Partial correlations investigating structure displayed significant negative correlations between TIV, ROI volume, and sequence learning for lobule V (*r* (104) = - 0.250, p_FDR_ =0.0425) and Crus I (*r* (104) = -0.231, p_FDR_ =0.0425), indicating lower lobule V and Crus I volumes were associated with increased learning performance (Figure 6).

**Figure 5.**
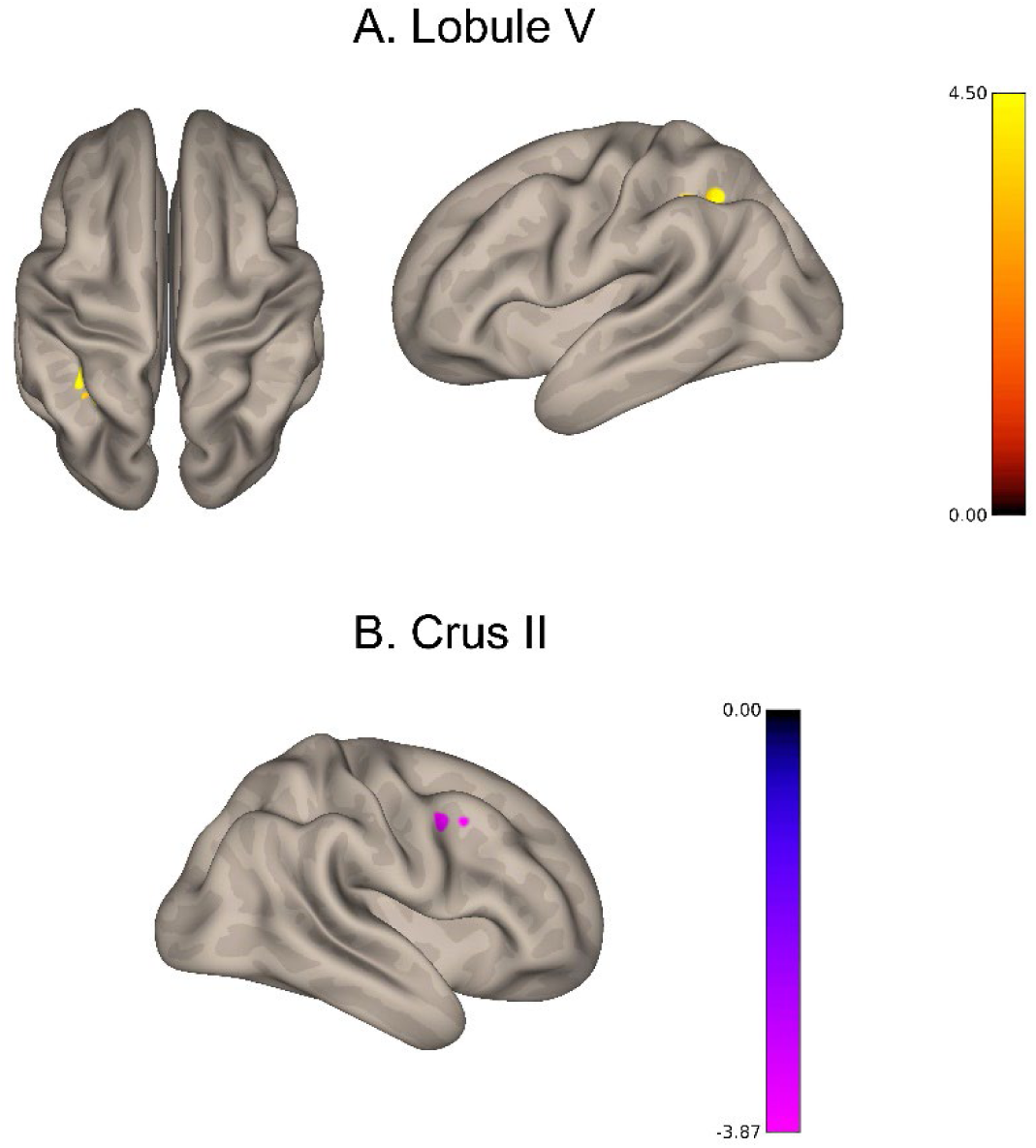
Bidirectional cortical functional connectivity (FC) is shown in ROIs with greater sequence learning. **A**. yellow represents greater FC between the inferior parietal gyrus and lobule V with greater sequence learning. **B**. purple displays lower FC between Crus II and the precentral gyrus with greater sequence learning.

**Figure 6.**
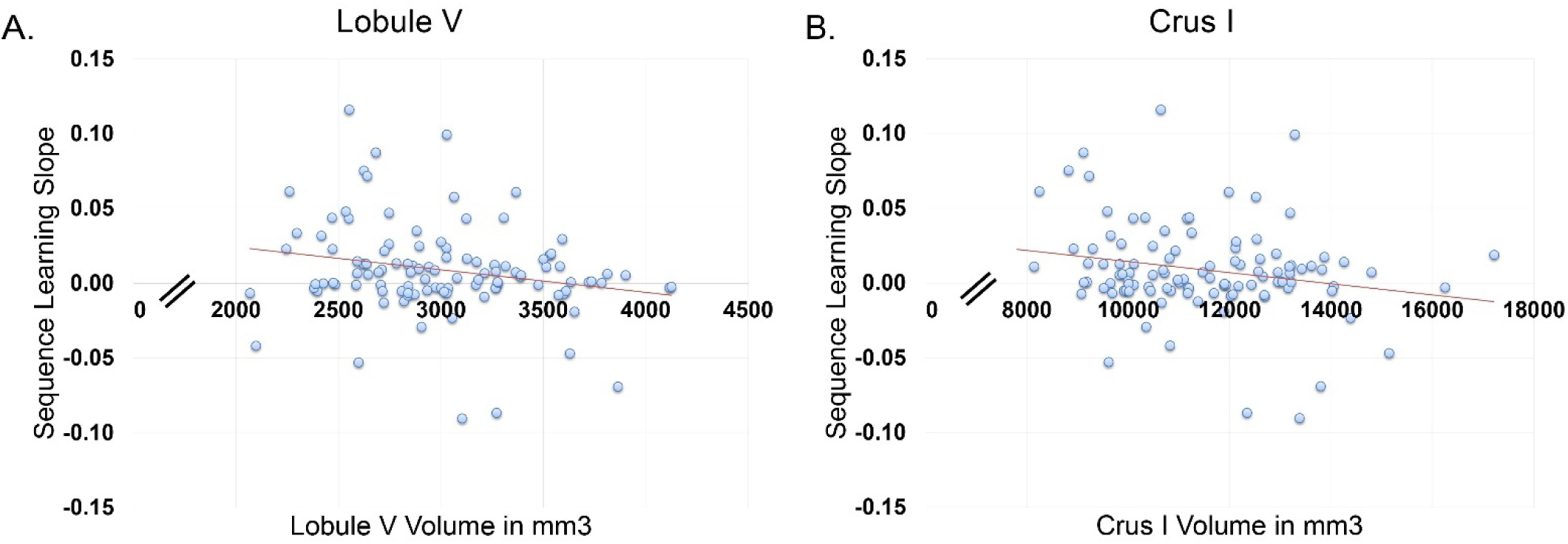
**A**. Lobule V volume shows a negative correlation with sequence learning. **B**. Crus I volume shows a negative correlation with sequence learning. These figures display rote correlations that do not account for TIV and are solely presented for visualization purposes.

**Table 6.** Coordinates of significant correlations between lobular cerebellar FC and Sequence learning.

| Functional Connectivity Correlations<br>with Sequence Learning Accuracy |  |  |  |  |  |  |
| --- | --- | --- | --- | --- | --- | --- |
| Region | X | Y | Z | Cluster<br>Size | T(105) | pFDR |
| **Crus II** |  |  |  |  |  |  |
| Precentral gyrus (BA 6) | 46 | 4 | 46 | 14 | -4.98 | 0.0004 |
| **Lobule V** |  |  |  |  |  |  |
| Inferior parietal gyrus (BA 40) | -34 | -42 | 36 | 20 | 5.90 | 0.0027 |
| Inferior parietal gyrus (BA 39) | -28 | -52 | 42 | 15 | 4.84 | 0.0078 |

Finally, we evaluated associations between the Purdue Pegboard peg assembly subtest separately from the motor composite to better understand relationships between a complex bimanual task requiring integrated cognitive and motor function and cerebellar metrics. Raw scores were converted to z-scores and a higher score indicates better performance on this task. Correlations between pegboard assembly and ROI connectivity were significant for Crus I, Crus II, and vermis VI (Figure 7; Table 7).

**Figure 7.**
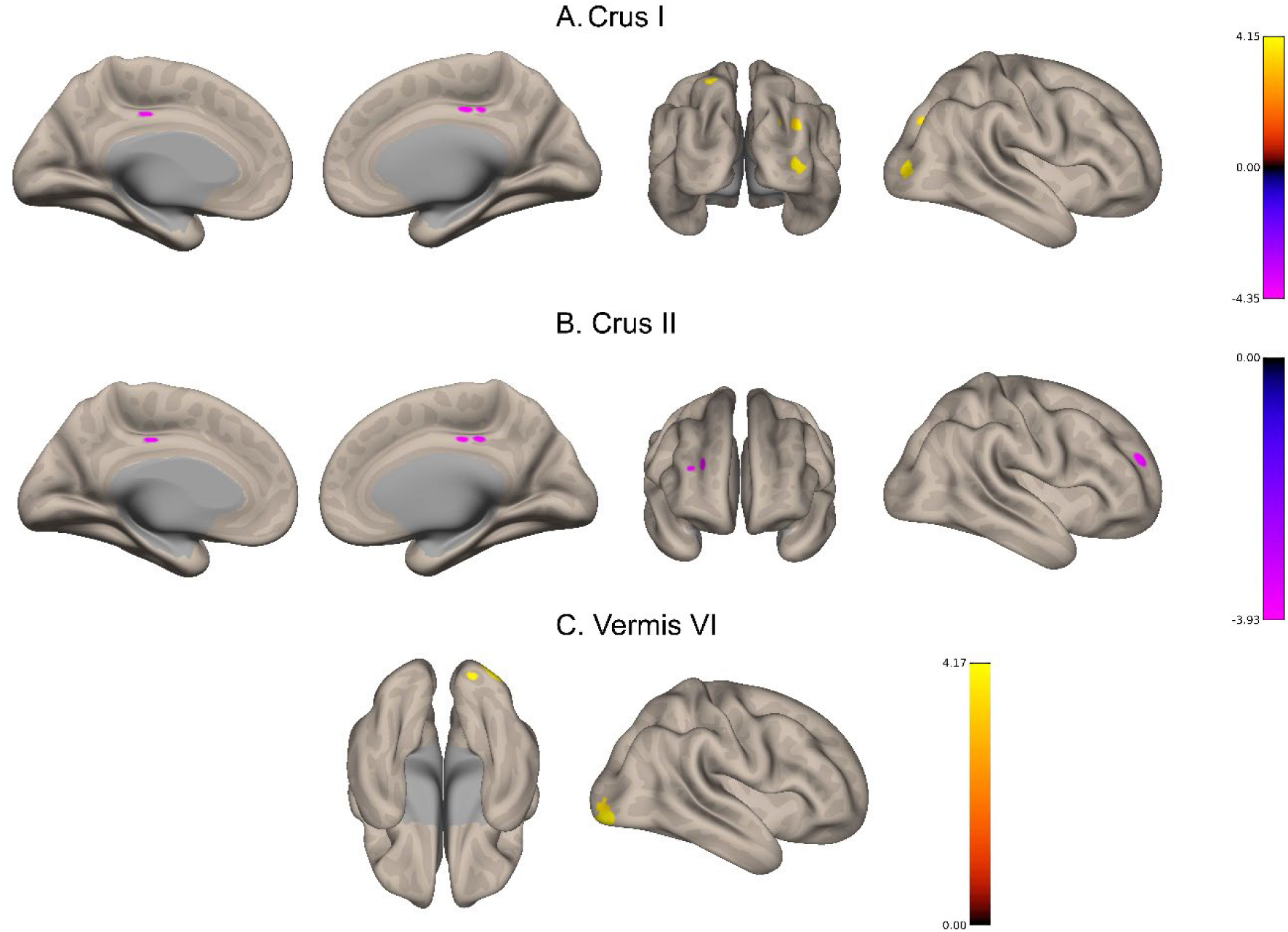
Bidirectional functional connectivity (FC) is shown in ROIs with better Pegboard Assembly performance. **A**. purple represents lower FC between Crus I and the mid-cingulate gyrus with better Pegboard Assembly performance; while yellow displays greater FC between Crus I and both the middle occipital and superior parietal gyri with greater Pegboard Assembly performance. **B**. purple displays lower FC between Crus II and both the middle frontal and mid-cingulate gyri with greater Pegboard Assembly performance. **C**. yellow represents greater FC between vermis VI and the inferior occipital gyrus with higher Pegboard Assembly performance.

**Table 7.** Coordinates showing significant correlations with lobular cerebellar FC and Pegboard Assembly.

| Functional Connectivity Correlations<br>with Pegboard Assembly |  |  |  |  |  |  |
| --- | --- | --- | --- | --- | --- | --- |
| Region | X | Y | Z | Cluster<br>Size | T(136) | pFDR |
| **Crus I** |  |  |  |  |  |  |
| Middle occipital gyrus | 32 | -78 | 32 | 29 | 5.12 | 0.0032 |
| Mid-cingulate gyrus | 2 | -18 | 42 | 20 | -6.27 | 0.0160 |
| Middle occipital gyrus | 36 | -78 | 2 | 19 | 4.94 | 0.0160 |
| Superior parietal gyrus | -22 | -58 | 62 | 15 | 5.10 | 0.4040 |
| **Crus II** |  |  |  |  |  |  |
| Middle frontal gyrus | 28 | 48 | 18 | 18 | -5.56 | 0.0347 |
| Mid-cingulate gyrus | 2 | -18 | 42 | 17 | -5.05 | 0.0347 |
| **Vermis VI** |  |  |  |  |  |  |
| Inferior Occipital gyrus | 32 | -86 | -8 | 95 | 6.64 | <0.0001 |

Specifically, better performance on pegboard assembly was correlated with greater connectivity between Crus I and regions of both the middle occipital gyrus and the superior parietal gyrus (Figure 7A). Interestingly, higher performance on pegboard assembly was also correlated with lower connectivity between Crus I and the mid-cingulate gyrus (Figure 7A). Better performance on pegboard assembly was significantly correlated with lower connectivity between Crus II and both the middle frontal gyrus and mid-cingulate gyrus (Figure 7B). Greater connectivity between vermis VI and the inferior occipital gyrus was significantly correlated with higher performance on pegboard assembly (Figure 7C). Partial correlations investigating lobular volume (controlling for TIV) demonstrated significant positive correlation between pegboard assembly performance and vermis VI (*r* (134) = 0.235, p_FDR_ =0.0300; Figure 8). Larger vermis VI volume was associated with better pegboard assembly performance. Partial correlations were not significant for the remaining ROIs (p_FDR_ > 0.05).

**Figure 8.**
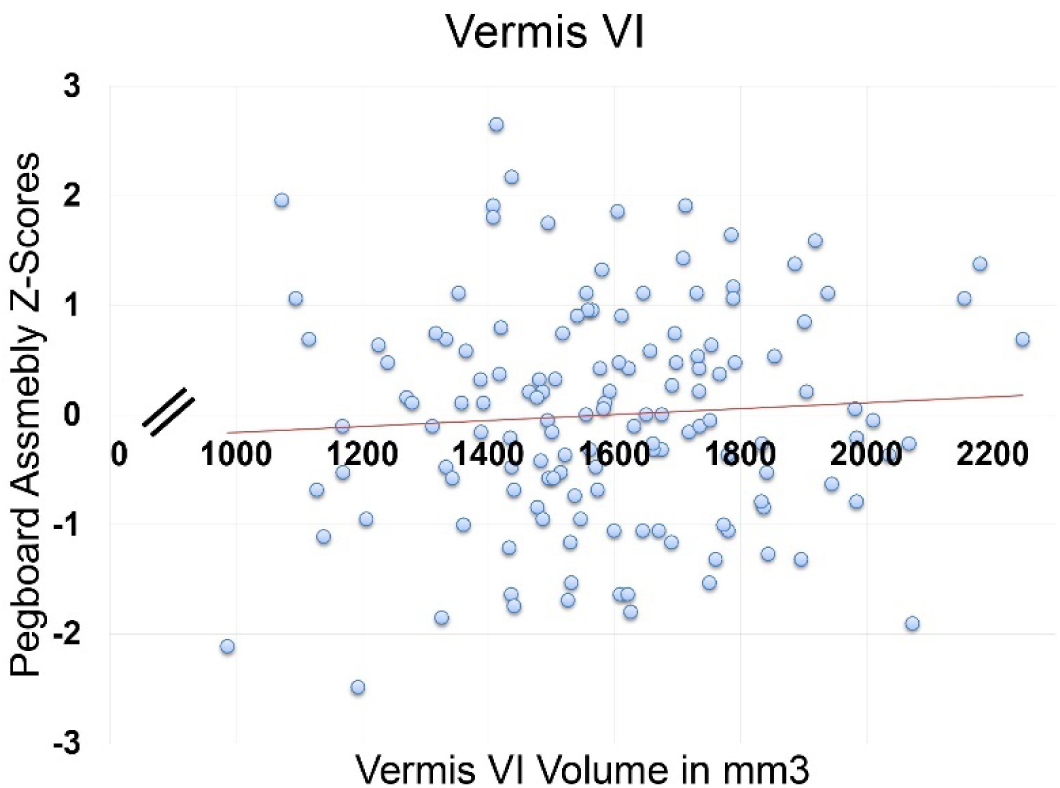
Larger vermis VI volume predicts better Pegboard Assembly performance. This figure displays rote correlations that do not account for TIV and are solely presented for visualization purposes.

The motor composite (all Purdue Pegboard subtests and bimanual grip strength) broadly measured upper limb motor abilities with a higher score indicating better motor performance. Motor composite correlations with lobular connectivity were significant for Crus II and vermis VI (Figure 9, Table 8). Specifically, lower connectivity between Crus II and the anterior cingulate gyrus was significantly correlated with better performance on the motor composite (Figure 9A). Interestingly, greater connectivity between vermis VI and occipital regions were correlated with better performance on the motor composite (Figure 9B). Partial correlations investigating regional volume demonstrated positive correlations between motor performance and lobules I-IV (*r* (134) = 0.244, p_FDR_ =0.0100), lobule V (*r* (134) = 0.308, p_FDR_ < 0.0001), Crus II (*r* (134) = 0.208, p_FDR_ =0.0188), and vermis VI (*r* (134) = 0.209, p_FDR_ =0.0188), indicating higher volumes were associated with better motor performance (Figure 10). Partial correlations were not significant for Crus I (p_FDR_ > 0.05).

**Figure 9.**
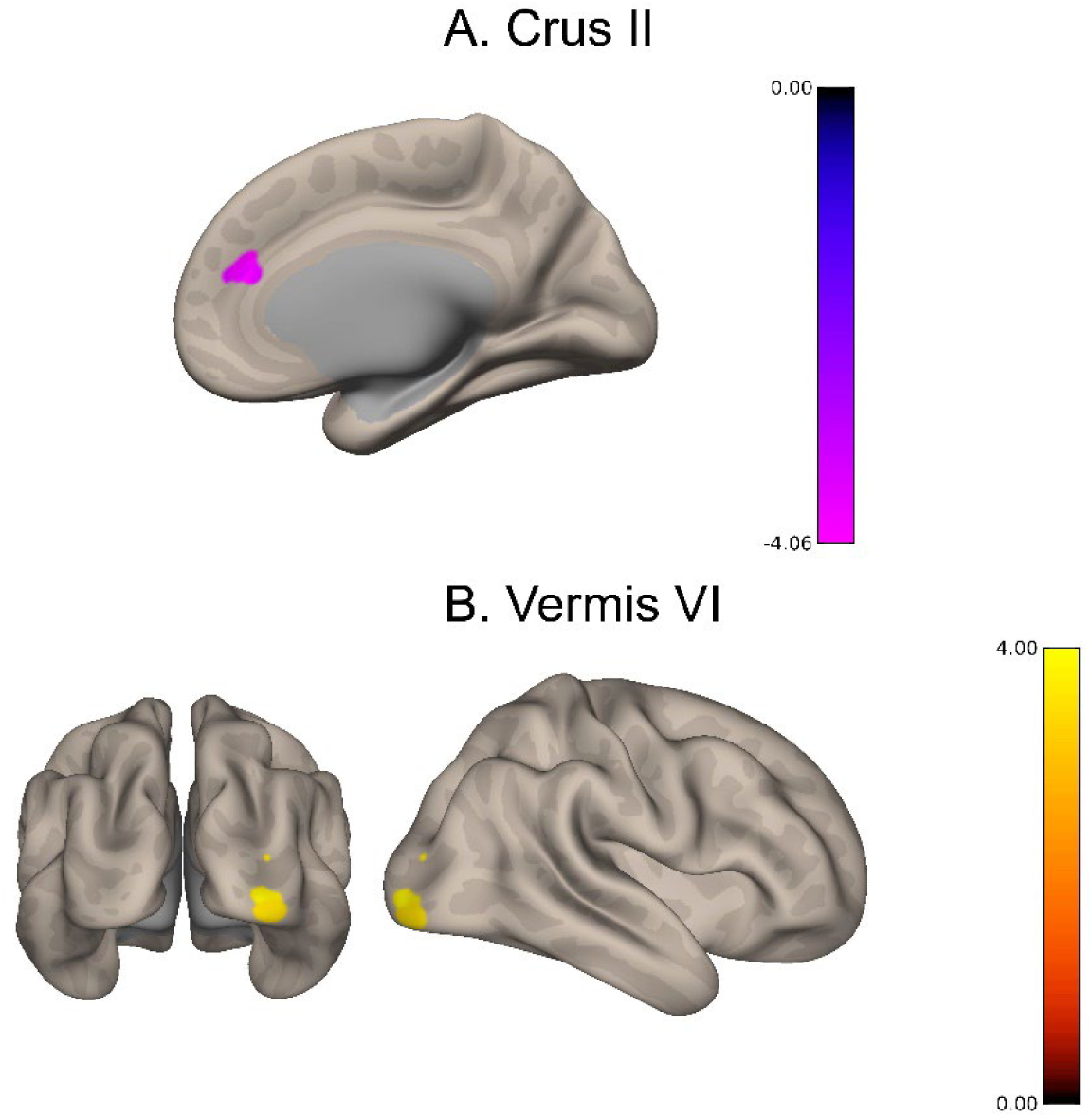
Bidirectional cortical functional connectivity (FC) is shown in ROIs with better performance across motor tasks. **A**. purple represents lower FC between Crus II and the anterior cingulate gyrus with better motor performance. **B**. yellow represents greater FC between Vermis VI and the lingual gyrus, middle occipital gyrus, and inferior occipital gyrus with better motor performance.

**Figure 10.**
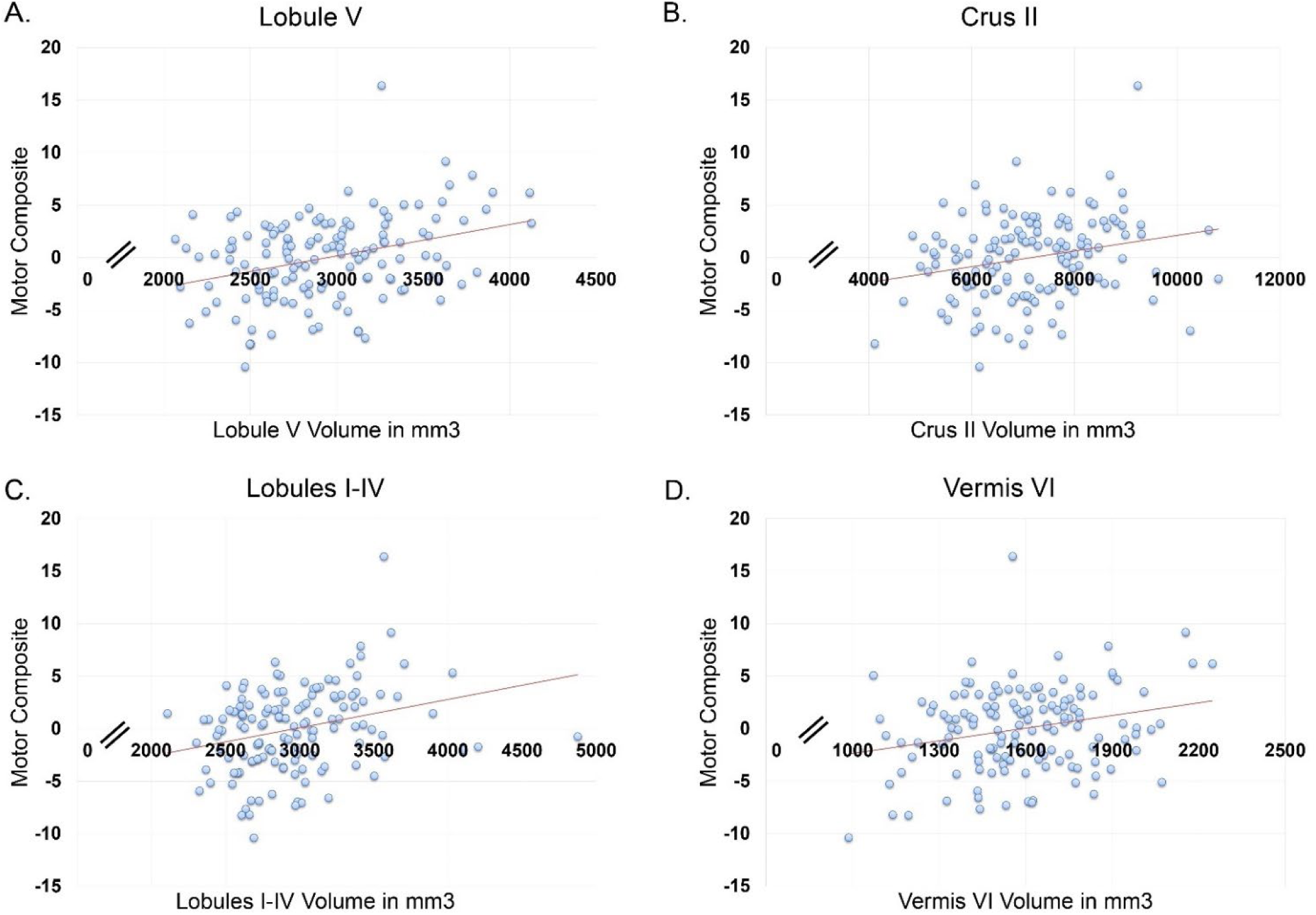
ROI Volume predicts higher performance across motor tasks. These figures display rote correlations that do not account for TIV and are solely presented for visualization purposes.

**Table 8.** Coordinates of regions showing significant correlations between lobular cerebellar FC and the motor composite score.

| Functional Connectivity Correlations<br>with Motor Composite |  |  |  |  |  |  |
| --- | --- | --- | --- | --- | --- | --- |
| Region | X | Y | Z | Cluster<br>Size | T(136) | pFDR |
| **Crus II** |  |  |  |  |  |  |
| Anterior cingulate gyrus | 10 | 42 | 20 | 28 | -5.64 | 0.0034 |
| **Vermis VI** |  |  |  |  |  |  |
| Lingual gyrus | 22 | -96 | -10 | 30 | 6.04 | 0.0016 |
| Middle occipital gyrus | 36 | -94 | 2 | 29 | 5.40 | 0.0016 |
| Inferior occipital gyrus | 32 | -88 | -8 | 20 | 4.45 | 0.0436 |

## Discussion

Here, we characterized relationships between age, cerebellar structure, and lobular connectivity across the adult lifespan. We further examined relationships between behavior and lobular cerebellar ROI structure and cortical FC in our healthy aging sample. Results revealed structural and functional associations with cross-sectional age as well as a variety of volumetric and cerebello-cortical connectivity relationships in lobules I-IV, V, Crus I, Crus II, and vermis VI. Notably, our study has the advantage of a relatively large sample size, allowing for robust statistical analyses. Additionally, we provided a comprehensive examination of behavior across multiple domains, coupled with detailed structural and resting state investigations at the lobular level. Overall, these investigations provide a more nuanced understanding of the cerebellum’s role in healthy aging, through the consideration of multiple behavioral domains, and the inclusion of individuals across the adult lifespan (age 35 and over).

### Age is Associated with Lobular Connectivity and Volume

The findings of this study exhibited both similarities and differences when compared to existing literature. Consistent with both our hypothesis and the results of previous studies examining cerebellar structure in aging, we found negative relationships between volume and greater age across ROIs when accounting for TIV (21,22). Two studies reported findings that were largely consistent with ours, demonstrating smaller relative volumes in lobules I-VI and Crus I with increased age; however, they did not find lower volumes in Crus II with aging (30,41). Cui and colleagues (20) found lower volumes in Crus I and Crus II in older as compared to young adults; however, this finding was confined to only 4 of the 12 cerebellar regions examined, in contrast to all 5 regions demonstrating a negative relationship between ROI volume and increased age in this study. Overall, our results suggest lower ROI volume across aging adults.

Functionally, our findings were mixed with respect to our hypotheses and consistency with the broader literature. We expected lower FC across cerebellar lobules to DMN regions with increased age. However, we found several areas of higher connectivity with increased age including lobules I-IV, Crus I, and Crus II with frontal, occipital, cerebellar, and more central regions of the brain (i.e., caudate and thalamus). While He and colleagues (26) examined cerebellar connectivity as a single cerebellar network, their findings showing intact cerebellar networks in aging are largely consistent with our results showing higher connectivity between these ROIs with increased age. Bernard and colleagues (27) found lower connectivity between both dorsal and ventral cerebellar dentate regions to the whole brain with increased age; however, as these regions are distinct from those investigated here, direct comparisons cannot be made. Broadly, higher connectivity between lobules I-IV, Crus I, and Crus II with regions such as the thalamus, anterior cingulum, and frontal regions may indicate relatively intact cerebello-thalamo-cortical pathways in healthy aging (71), that may relate to healthy aging outcomes.

Consistent with past work on connectivity in aging more generally, we saw relationships with age that were bidirectional. Lower connectivity was seen between Crus II and lobule V with temporal, parietal, occipital and cerebellar regions with increased age. In line with our hypothesis, we found lower connectivity in the precuneus, lateral parietal, and medial temporal regions which are included in the DMN (72). Crus II specifically has been theorized to play a role in the DMN (28). These results may indicate anatomically specific cerebellar involvement in the DMN. Further, this may indicate cerebellar circuits that are more vulnerable to aging. As these relationships have been less frequently explored at the lobular level, our findings may provide more nuance to the mixed findings in the literature (26,27,29). That is, higher connectivity between lobules I-IV, Crus I, and Crus II and frontal, occipital, cerebellar, and more central regions of the brain (i.e., caudate and thalamus) could indicate intact networks in aging; whereas, lower connectivity between Crus II, lobule V and temporal, parietal, occipital and cerebellar regions may imply vulnerable networks in aging.

### Behavior is Associated with Lobular Connectivity and Volume

We examined a variety of cognitive variables to assess cerebello-cortical functional relationships and volumetric associations with cognition. As predicted, larger relative Crus I and vermis VI volumes correlated with better performance on our first cognitive composite variable (Comp1) that broadly assessed attention, processing speed, and working memory on paper and pencil tasks. Similarly, Bernard and Seidler found greater Crus I volume was linked to better performance on a higher-level cognitive processing task of choice reaction time (30). Taken together, this suggests that relative Crus I volume may be useful in predicting cognitive performance for tasks involving processing speed, attention, high-level processing, and working memory. Vermis VI volume might also be an important region to investigate further for a role in cognition. Notably, cortical FC correlates for our lobular ROIs of interest to Comp1 were not significant. This was counter to expectations given functional imaging work implicating these cerebellar regions in working memory and attention in young adults (37,38).

Our second cognitive composite (Comp2) broadly represents episodic memory, executive function, and motor learning on computer-based tasks, and we saw distinct results. Consistent with our hypothesis, we demonstrated a significant positive correlation between Comp2 performance and connectivity between Crus I and the superior parietal gyrus. The superior parietal gyrus has been linked to spatial shifting and attention (73). These results may be related to the spatial attentional demands required in the Stroop and sequencing tasks. The correlational relationship seen between Crus I and the superior parietal gyrus with better performance on Comp2 may also provide support for successful cognitive modulation via the cerebello-thalamo-cortical pathway which extends to parietal cortices (71,74–76). It is notable that we did not find a positive relationship between Crus II connectivity and Comp2 as Crus II connectivity to the precuneus and posterior cingulate has previously been linked to working memory performance (28). It is possible that the combination of cognitive measures for Comp2 was too broad to involve this lobular region. Comp 2 also did not correlate with volume across remaining ROIs, contrary to our initial predictions.

In terms of the Stroop effect for reaction time, a stronger Stroop effect (slower performance on incongruent items relative to performance speed on congruent items) was correlated to higher connectivity between lobule V and the postcentral gyrus. The postcentral gyrus is part of the somatosensory network (77), which is less often related to cognitive tasks. A stronger Stroop effect was also correlated with lower connectivity between Crus II and lobule VI, between lobule V and both the middle frontal gyrus and inferior frontal gyrus, and between vermis VI and the middle frontal gyrus. Thus, slowed performance on more complex items requiring executive function abilities was linked to lower connectivity between Crus II, lobule V, and vermis VI and both frontal and cerebellar regions. Broadly, the aforementioned frontal regions have displayed involvement in networks (i.e., DMN and frontoparietal) which show lower connectivity with increased age (6,26,49,50). These results may suggest cerebellar involvement in DMN and frontoparietal networks as they relate to executive function, highlighting these connections as regions of vulnerability for executive function performance with increased age. Our results were consistent with the cerebellar functional topography, which linked lobule V to motor planning and attention and Crus II to interference resolution (37), domains necessary to perform well on the Stroop task.

The Purdue Pegboard peg assembly subtest was used to evaluate correlational relationships between cerebellar activation and a task of integrated cognitive and motor function. Better performance on this task was linked to higher connectivity between Crus I and occipital and parietal regions as well as vermis VI and the inferior occipital gyrus.

In Stoodley and colleagues’ functional topography study (46), overt movement was linked to sensorimotor cortices and lobules IV-V; whereas cognitive tasks engaged prefrontal and parietal cortices. The pattern of results shown here are somewhat consistent with Stoodley and colleagues’ findings but may more specifically represent networks or circuits that represent complex cognitive-motor integration. Consistent with this notion, better performance on peg assembly was also associated with lower connectivity between Crus I and the cingulate gyrus as well as Crus II and both frontal and cingulate regions. Regarding lobular volumes, there were positive correlations between peg assembly performance and vermis VI volume. Functional activation in vermis VI has been linked to motor tasks in both young and older adults as well as working memory in older adults (25). However, complex cognitive-motor tasks have yet to be associated with vermal volume, especially in older adults.

Motor composite correlations exhibited bidirectional FC relationships. We predicted that higher FC between lobules I-IV and V to motor association regions would display positive relationships with better motor performance. We found that better motor performance was correlated with higher connectivity between vermis VI and occipital regions; however, connectivity was lower between Crus II and the mid-cingulate gyrus. The occipital and mid-cingulate gyrus regions are association areas which have been found to correlated with cerebellar connectivity (47), especially in the context of distinct motor and cognitive functions (52). Cerebellar lobules have exhibited distinct relationships with motor cortical, cognitive, and association regions (52). Surprisingly however, these relationships were not seen with lobule V which has been extensively linked to motor function and networks (37,39,46). In contrast, structural findings did display positive relationships between the motor composite and volume across all ROIs, consistent with Koppelmans and colleagues (41) findings showing links between Crus I and lobules I-VI with motor behavior. Thus, we exhibited robust structural findings for ROI volume as it related to motor function in aging adults.

Together, our results shed light on both structural and functional relationships with age, cognition, and motor function. We found lower volumes in lobules I-IV, V, Crus I, Crus II, and vermis VI with increased age as well as bidirectional cerebellar connectivity relationships with increased age, consistent with literature on FC in aging (26–28,49). Further, we revealed unique associations in both cerebellar structure and connectivity with comprehensive behavioral measures in a healthy aging population. However, there are also some limitations to consider in interpreting this work. First, our data were cross-sectional in nature which does not allow for causal inferences or an understanding of individual changes over time. Further, all behavioral tasks were administered outside the scanner. Therefore, correlational relationships between behavior and brain measures should be interpreted with caution as the data were collected across several sessions.

## Conclusion

This study characterized relationships between age, cerebellar structure, and lobular connectivity across the adult lifespan. Structural and functional associations were shown with cross-sectional age. Cerebello-cortical connectivity was demonstrated across ROIs in the context of age and/or behavior. These results replicate and extend findings in the literature supporting a role for the cerebellum in behavior in advanced age. Future studies can build on these data for clinical application and translational value as they provide a strong baseline with respect to the broad age range and diverse task battery employed here.

## Acknowledgment

This work was supported by R01AG065010 to J.A.B. This work was further supported by the Texas Virtual Data Library (ViDaL), a high-performance cluster, funded by the Texas A&M University Research Development fund. In this cluster the imaging analyses for the current work were carried out using the resources provided by the Texas A&M High Performance Research Computing organization.

## Notes

### Competing Interest Statement

The authors have declared no competing interest.

